# Isolation of Secretory Immunoglobulin A (sIgA) from nasal secretions collected in saline and Viral Transport Medium

**DOI:** 10.1101/2025.06.26.661459

**Authors:** Rohit Bhat, Bhakti Surve, Sunil Nadkarni, Rita Mulherkar

## Abstract

Secretory immunoglobulin A (sIgA) is the major immunoglobulin found on all mucosal surfaces, including the nasal cavity and the lining of the gut. It is the first line of defence against pathogens which enter through the nasal cavity or mouth. sIgA1 binds to a lectin isolated from the tropical jackfruit, known as jacalin. Jacalin binds specifically to the D-galactose moiety of sIgA1 and not to sIgA2 which has different sugars. Here we describe a simple and inexpensive method to isolate jacalin from jackfruit seeds, prepare jacalin-Sepharose conjugated beads and isolate sIgA1 from nasal secretion collected in saline as well as in Viral Transport Medium (VTM). The sIgA identity was confirmed by Western blotting using specific antibodies against sIgA α heavy chain. Our approach has potential implications as studying sIgA in nasal secretions provides a direct measure of the effectiveness of nasal vaccines at the site of pathogen entry, making it a critical biomarker for both vaccine development and evaluation.

This research did not receive any specific grant from funding agencies in the public, commercial, or not-for-profit sectors.

## Introduction

Mucous membrane covers a huge area along the epithelial lining of the alimentary canal, respiratory tract, urinogenital tissue as well as conjunctiva covering the eyes, which is a major barrier protecting the body against pathogenic organisms, allergens and carcinogens (1-3). The predominant antibody protecting the mucous membrane lining in the mucosal tissue is secretory IgA1 (2, 4, 5). sIgA constitutes greater than 80% of all antibodies produced in mucosa-associated lymphoid tissues in humans. It is the first line of defence against any inhaled, ingested or sexually transmitted pathogen (6). It is present in all body secretions such as saliva, tears, nasal secretions, gastric juices as well as milk. It is also the predominant antibody present in human milk (7). sIgA binds pathogens via its glycoproteins and neutralizes the catalytic activity of the microbial enzymes, thus preventing their entry in the gut (6, 8). Also, since large number of sIgA are cross-reactive, the number of antibodies required to inactivate the pathogen are fewer in number. sIgA carry the gut associated antigens to dendritic cells in the gut associated lymphoid tissue for immune priming (6).

Respiratory viruses such as SARS-CoV2 enter through the nasal cavity and oral route. When evaluating nasal vaccines, especially those targeting respiratory pathogens (e.g., influenza, RSV, SARS-CoV-2), measuring sIgA levels in nasal secretions could help assess whether the vaccine successfully stimulates mucosal immunity (9). Also, it indicates the strength and duration of the local immune response. This may correlate with protection better than systemic antibodies such as IgG, for mucosal infections (10).

We present a simple method to isolate sIgA from nasal secretions collected in saline as well as in Viral Transport Medium (VTM). VTM is routinely used to safely transport test samples to a laboratory for testing. Our approach has potential implications not only in the understanding of mucosal immunity but also in developing intranasal vaccines that can induce sIgA. Understanding the immune response is of paramount importance.

## Methodology

Institute Ethics Committee approval was obtained before collecting biofluid from the individual. Informed consent was taken from all subjects.

### Purification of jacalin

Jackfruit (*Artocarpus integrifolia*) seeds were taken for extraction of lectin jacalin. It was further purified using affinity chromatography on affinity chromatography on cross-linked guar gum, a polysaccharide comprising galactomannan matrix as described earlier (11).

About 100 grams of jack fruit seeds were homogenized in a blender and the fine powder was air dried at room temperature. Each of the subsequent purification steps was carried out at 4°C unless mentioned otherwise. The dry powder was soaked in 20 mM phosphate buffer containing 150 mM NaCl and 0.02% sodium azide (PBS, 1: 10; w/v) and was stirred overnight. The extract was passed through a cheese cloth and the filtrate was centrifuged at 10000 rpm at for 30 minutes. To the supernatant ammonium sulphate was added up to 70% saturation, stirred for 30 minutes and kept overnight. Then it was centrifuged at 8000 rpm for 30 minutes and the pellet was collected and dissolved in minimum volume of PBS and dialysed extensively against PBS. The dialysate was again centrifuged at 8000 rpm for 30 minutes and the protein in the supernatant estimated. Ten to 15 ml of dialysate was passed through guar-gum column (2.5 × 30 cm) pre-equilibrated with PBS. The breakthrough was reloaded in order to ensure complete binding of jacalin to the matrix. The column was then washed with PBS until the OD_280_ fell below ∼ 0.01. The bound protein was eluted with 0.2 M galactose at room temperature. The purity, homogeneity and estimation of the purified protein was assessed by SDS-PAGE.

### Preparation of jacalin-coupled Sepharose 6B affinity matrix

Jacalin was coupled to Sepharose 6B, activated by divinyl sulfone, essentially as described earlier (12). In brief, 20 ml of packed Sepharose 6B was thoroughly washed with distilled water. To activate the gel, about 20 ml of 0.5 M carbonate/bicarbonate buffer, pH 11, was added to the wet gel followed by the addition of 2 ml of divinyl sulfone. The mixture was kept at room temperature for 70 min under continuous stirring and was subsequently transferred to a glass sintered funnel and carefully washed with distilled water. The activated gel was then suspended in 20 ml of 0.5 M carbonate/bicarbonate buffer, pH 10, containing 200 mg of purified jacalin and the coupling reaction was allowed to proceed for 24 hours at 4 °C. The extent of coupling was determined by measuring the absorbance of the solution obtained from the coupling mixture after passing through the sintered funnel as well as by estimating the protein content. The coupled gel was extensively washed with water and suspended in carbonate/bicarbonate buffer, pH 8.5. Then 200 µL of 2-mercaptoethanol was added to the suspension and mixed for 3 hours. The gel was finally washed with water and suspended in equilibrating buffer containing 0.02% azide till further use. Just before use the gel was equilibrated in PBS, pH 7.6 and packed in a 26×1cm column with PBS as the wash buffer.

### Biofluid and Affinity column chromatography

Institute Ethics Committee approval was obtained before collecting biofluid from the individual. Milk was collected from healthy mothers, after taking informed consent. Five ml of milk was collected and kept frozen at -20°C until further use. Milk was thawed by warming in a 37°C and then spun at 100,500xg at 4°C. The fat layer was removed and clear skimmed milk was loaded on the jacalin affinity column and allowed to pass through the gel. The column was washed with PBS and 2 ml fractions collected until the absorbance at 280 nm of the effluent reached baseline. The bound SIgA was eluted with 0.1M Melibiose and used as a positive control. Absorbance at 280 nm of all fractions was taken on a Shimadzu Spectrophotometer. Protein fractions eluted with Melibiose were saved and stored at 4°C until further use.

### Isolation of SIgA from nasal secretion

Nasal secretion was collected from consenting participants, with a moistened cotton swab by inserting it in both the nostrils and swirling it inside the nostril for a few seconds. The swab was immediately placed in a tube containing 2 ml saline. For collecting sample from nasopharyngeal region, a minitip swab with flexible shaft was used and transferred in the Viral Transport Medium (VTM) containing tube. The tubes were labelled and kept at -20°C until further use. Protein estimation in the samples was done using BCA method. To isolate sIgA, the tube was thawed and 1 to 1.5ml of the saline / VTM solution with the nasal secretion was taken in a microfuge tube. Hundred microlitres of jacalin-Sepharose6B gel suspension was added to the tube and placed on a shaker for 1 hr at RT. The gel was washed thrice by centrifuging the tube in a micro spin for 5 min at 10000 rpm. Finally, bound sIgA was eluted with 100ul Melibiose sugar twice by placing on a shaker for 20 min at RT. The eluted SIgA was collected in a separate tube and concentrated on a Centricon tube (mw cutoff 3,000 Da, Millipore Sigma). The membrane was washed twice with distilled water and eluted from the membrane in a minimum amount of water. The protein concentration in the eluted sample was quantitated.

### SDS-PAGE and Western blotting

SDS-polyacrylamide gel electrophoresis (SDS-PAGE) was carried out on 12% SDS-PAGE slab gel in the buffer system of Laemmli (13). Thirty microlitres of sample from unbound proteins and eluted proteins were run on the gel under non-reducing and reducing conditions. The gel was stained with Coomassie Brilliant Blue or silver stained. Proteins from an identical gel were transferred to a PVDF membrane. Blocking was carried out for one hour in 5% milk powder, in Tris-buffered saline containing 0.1% Tween 20 (TBST) at room temperature. The blot was incubated with biotinylated goat anti-human α chain of sIgA (5 µg/ml, Mabtech, Sweden) for 2 hr at RT. After the washing the membrane several times with PBST, the blot was incubated in HRP-Streptavidin (1:1000 dilution) for 2 hr at RT, rinsed with TBS and developed with freshly made DAB (1mg/ml in TBS) containing H_2_O_2_.

## Results

In order to purify sIgA on affinity chromatography, we first isolated jacalin, an sIgA binding lectin from different batches of jackfruit seeds. Jackfruit seed extract was loaded on a Guar gum column and the bound jacalin was eluted with D-galactose. The purity of the eluted jacalin was checked on a 12% SDS-PAGE subjected to silver staining (Fig 1). Jacalin is a tetramer made up of two 16kDa peptides and two 14kDa peptides as seen in the Fig. 1.

**Fig. 1.**
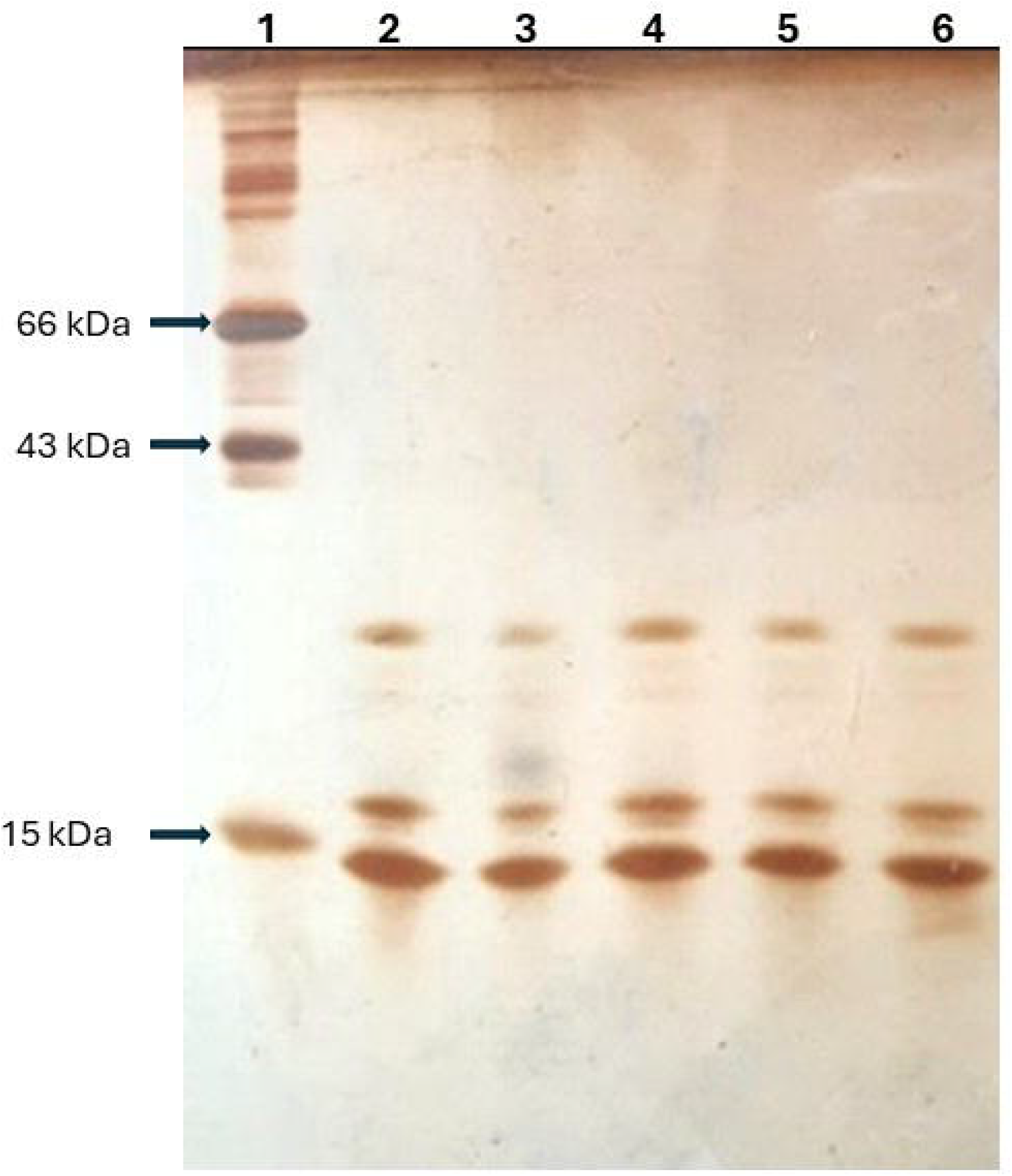
Different batches of purified jacalin were run on 12% SDS-PAGE and subjected to silver staining. Lane 1: Molecular weight markers; Lane 2 to 6: Different batches of purified jacalin.

Further, for affinity chromatography, the purified jacalin was conjugated to Sepharose 6B beads. Initially sIgA from defatted milk was isolated on the jacalin-Sepharose column to confirm its use to purify sIgA. Purified sIgA from milk gave the expected 3 bands under reducing conditions – Secretory Component (75kDa), Heavy chain (55kDa) and light chain (25kDa, Fig. 2A).

**Fig. 2.**
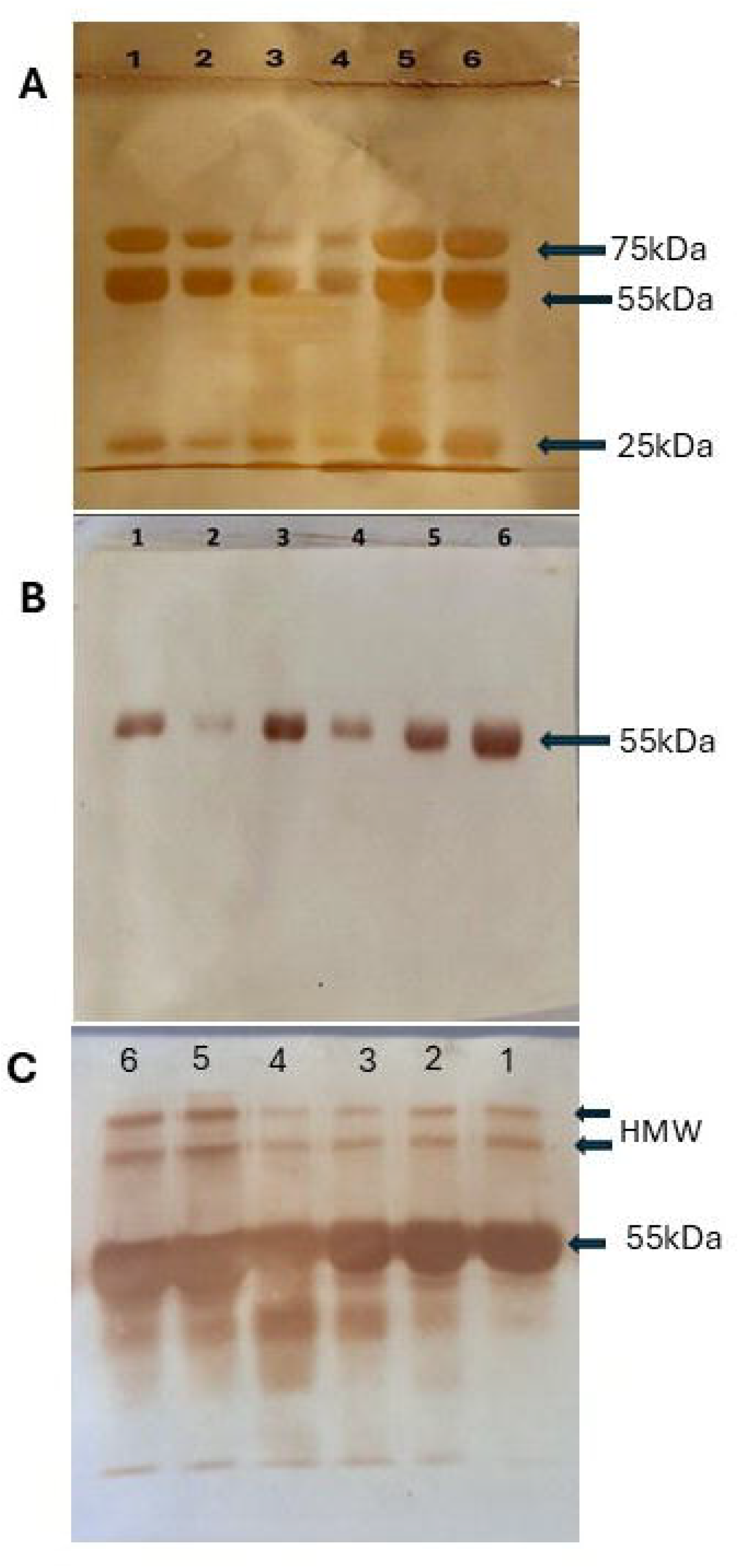
Protein from nasal secretions collected in different media were subjected to Western blot analysis. (A) Silver-stained gel; (B) Western blot of proteins (low concentration) from nasal secretions probed with anti-α heavy chain of sIgA; (C) Western blot of proteins (high concentration) from nasal secretions probed with anti-α heavy chain of sIgA. Lanes 1&2: sIgA from nasal secretions in saline; Lanes 3&4: sIgA from nasal secretions in VTM; Lanes 5&6: sIgA from defatted milk.

In order to isolate very small quantities of sIgA from the nasal secretions collected via moistened nasal swabs, the jacalin-Sepharose beads were added to the nasal secretions collected either in saline or VTM, in microcentrifuge tubes. The bound sIgA was eluted with the sugar Melibiose and the salts removed by spinning on a Centricon tube. Aliquots of the eluted sIgA from saline swabs, VTM swabs and defatted milk (control) were run on two 12% SDS-PAGE. One gel was silver stained (Fig. 2A) and the other gel was transblotted to a membrane and probed with biotinylated anti-sIgA α heavy chain, developed with HRP-DAB. A band corresponding to α heavy chain (∼55kDa) was seen (Fig. 2 B, C). When higher protein concentrations were run two high molecular weight bands (HMW) seen in the Western blot (Fig. 2C). These could be the multimeric and dimeric forms of sIgA detected by the sIgA specific antibodies as reported by Chen et al (9).

## Discussion

The mucosal lining in humans is exposed to numerous pathogenic as well as symbiotic microorganisms (6). The mucosal B lymphocytes present in lymphoid organs, play an important role in immunity as they differentiate into antibody secreting cells and produce large amounts of IgA in the mucosa. sIgA constitutes greater than 80% of all antibodies produced in mucosa-associated lymphoid tissues in humans. Human milk is also known to contain sIgA as one of the major proteins (8). Human colostrum has high concentrations of sIgA suggesting an important role in passive immune protection against infections in the babies (7). Passive administration of human sIgA in a mouse model of infection with M. tuberculosis could inhibit the infective potential of the pathogen (7).

Most of the infections involve the mucosae, but development of vaccines against these pathogens is still lacking. There are reports that mucosal immunization elicits immune response at local and distal mucosal sites as well as systemic immune response (1, 10). Intranasal vaccines can elicit a sIgA response to build a first line immune barrier against the pathogens and induce broadly neutralizing antibodies (9, 10). In order to shed more light on mucosal immunity, sIgA has been isolated from the nasal secretions from the vaccinated individuals in a non-invasive manner (10). sIgA can also be used as delivery vectors to deliver vaccines in the mucosa.

Secretory Immunoglobin A is involved in both protection against and pathogenesis of various diseases (14). Altered levels of sIgA are associated with immune dysregulation. Therefore, it can be used a marker for IgA nephropathy, celiac diseases, Crohn’s diseases, etc. Salivary sIgA plays a major role in oral health including in immune compromised cancer patients (15). Low sIgA is saliva is associated with increased risk of dental caries (16). Low levels of sIgA predispose individuals to recurrent mucosal infections including allergies and autoimmune diseases. Salivary sIgA has been used as a predictor of Upper Respiratory Tract Infections in elite rugby union players (17). Monitoring sIgA may provide an assessment of a team-sport athletes risk status for developing upper respiratory tract symptoms (18). Parasitic infestation of the gut also is known to elicit a nasal sIgA response. A nasal swab may prove a more amenable test in screening for parasitic infestation as compared to stool examination. There is some evidence to say that serum sIgA can correlate with systemic manifestations of disease. A simple method for estimating nasal sIgA may open the avenue for further work on a non-invasive test to calibrate and choose medications in conditions in auto-inflammatory diseases (19), such as sero negative spondyloarthropathy, Rheumatoid arthritis and inflammatory bowel disease.

We have optimized a cheap and simple method for purifying jacalin, an sIgA binding lectin, from jackfruit seeds, preparing affinity column with jacalin-Sepharose 6B beads, and using it to purify sIgA from milk as well as from nasal secretions. Chen et al (9) have reported collection of nasal secretions by nasal irrigation device for nasal antibodies purification. Here we describe a non-invasive and simple method for purifying sIgA using in-house prepared jacalin affinity Sepharose beads.

During the COVID pandemic, nasal swabs were collected in Viral Transport Medium to inactivate the virus and check for presence of the virus. We collected the left-over VTM containing the nasal secretions and isolated sIgA from VTM. We assume that this would be an ideal source to study sIgA in virus-infected patients and understand the immunology better. To the best of our knowledge, sIgA has not been purified from nasal secretions from patients, collected in VTM.

## Abbreviations

sIgA: Secretory Immunoglobin A
VTM: Viral Transport Medium
SDS-PAGE: sodium dodecyl sulphate-polyacrylamide gel electrophoresis
HRP: Horse Radish Peroxidase
DAB: 3,3’-diaminobenzidine.

## Acknowledgements

We wish to thank Shi Vithalrao Joshi Charities Trust for supporting our work. We also thank Virology Department at BKL Walawalkar Hospital, Sawarde for providing the VTM samples. All individuals who willingly provided nasal and milk samples are also acknowledged.

